# A molecular toolkit for heterologous protein secretion across Bacteroides species

**DOI:** 10.1101/2023.12.14.571725

**Authors:** Yu-Hsuan Yeh, Vince W. Kelly, Rahman Rahman Pour, Shannon J. Sirk

## Abstract

Bacteroides species are abundant and prevalent stably colonizing members of the human gut microbiota, making them a promising chassis for developing long-term interventions for chronic diseases. Engineering these bacteria as on-site production and delivery vehicles for biologic drugs or diagnostics, however, requires efficient heterologous protein secretion tools, which are currently lacking. To address this limitation, we systematically investigated methods to enable heterologous protein secretion in Bacteroides using both endogenous and exogenous secretion systems. Here, we report a collection of secretion carriers that can export functional proteins across multiple Bacteroides species at high titers. To understand the mechanistic drivers of Bacteroides secretion, we characterized signal peptide sequence features as well as post-secretion extracellular fate and cargo size limit of protein cargo. To increase titers and enable flexible control of protein secretion, we developed a strong, self-contained, inducible expression circuit. Finally, we validated the functionality of our secretion carriers in vivo in a mouse model. This toolkit should enable expanded development of long-term living therapeutic interventions for chronic gastrointestinal disease.

## Introduction

Engineered living therapeutics comprise an emerging class of microbial cell-based treatment strategies utilizing both native and engineered strains to function as biological machines that mediate disease prevention and treatment from within the human body. Achievements in this field are driven by a continuous expansion of our knowledge of host-microbiota interactions^1^ combined with ongoing innovations in synthetic biology tools^2^. These advances have supported numerous studies demonstrating that commensal gut bacterial species can be engineered to perform metabolic functions^3–6^ or produce and deliver therapeutic compounds from within the gastrointestinal (GI) tract^4,7–11^. Notably, ∼90% of the bacterial species in the average human gut belong to only two phyla – *Bacteroidetes* and *Firmicutes*^12^– and ∼15% of the entire gut microbial community belongs to a single dominant genus: *Bacteroides*^13,14^. Furthermore, a longitudinal study of human gut microbiota stability of healthy adults in the United States showed that the colonization of *Bacteroides* species can remain stable for up to 5 years^15^. Together, these features potentiate the use of *Bacteroides* species as a robust chassis for developing living therapeutics to achieve *in situ*, long-term treatment and monitoring of chronic diseases such as cancer, diabetes, and inflammatory bowel diseases (IBD). Critically, recent extensive studies of *Bacteroides* species provide a powerful selection of tools for genetic manipulation^16–24^, biocontainment^25^, and tunable engraftment^26^, further advancing their application from bench to bedside.

When engineering bacteria for therapeutic or diagnostic purposes in which protein-based products must function in the extracellular space, the ability of the microbial chassis to secrete heterologous cargoes is a key selection criterion^27,28^. While the Bacteroides genus represents an attractive collection of target species for this purpose, all Bacteroides species are Gram-negative, which presents technical challenges for efficient protein secretion. Unlike Gram-positive bacteria, which have a single lipid membrane and readily secrete heterologous cargoes outside of the cell through both the general secretion pathway (Sec) and twin-arginine translocation (Tat) pathway as long as the target protein is fused to an appropriate signal peptide (SP)^29,30^, protein secretion from double-membraned Gram-negative bacteria is more complex and requires additional cellular machinery^31^. Thus far, eleven different secretion systems have been identified in Gram-negative bacteria, referred to as the type 1 secretion system (T1SS) through T11SS^31,32^. Some of these secretion systems are well characterized and widely used for heterologous protein secretion in non-native hosts, such as T1SS^33^ and T3SS^34–36^. However, these secretion systems are either poorly conserved or completely absent from all Bacteroides species studied to date^37,38^. A few studies have reported that fusing a therapeutic protein with SPs from Bacteroides fragilis enterotoxin or OmpA (BT_3852) of Bacteroides thetaiotaomicron (B. theta)^16,39–41^ yields detectable levels of secretion; however, the highest reported titers of ∼900 pg/mL is still orders of magnitude lower than the secretion titers observed with other well-studied Gram-negative and Gram-positive bacteria, often exceeding 100 µg/mL^29,42^.

To address these limitations and capitalize on the potential of *Bacteroides* species as a powerful platform for developing engineered living therapeutics, we systematically investigated the heterologous protein secretion capability of both endogenous and exogenous secretion systems in *B. theta.* Our investigations reveal a suite of full-length proteins and lipoprotein SPs derived from native *B. theta* secretory proteins that can deliver functional antibody fragments and reporter proteins into the extracellular space. We show that these secretion carriers are broadly functional across multiple *Bacteroides* species. We also define the sequence features and preferred amino acid composition of lipoprotein SP that can drive high-level secretion. To further refine the usage of secretion carriers, we evaluated the post-secretion fate of protein cargoes exported via full-length fusion partners and lipoprotein SPs and observed both OMV-dependent and OMV-independent secretion. By selecting specific secretion carriers from our collection, we are able to direct secreted proteins to specific target destinations: freely soluble in the extracellular space; bound to the external surface of OMVs; or held within the OMV lumen. Additionally, toward full characterization and optimization our system, we identified the cargo size limit for lipoprotein SP-mediated secretion of heterologous proteins and also developed a strong, self-contained, inducible system for driving robust protein expression in *Bacteroides* species. Finally, we validated the activity of our secretion carrier platform *in vivo* using *B. theta* engineered to express and secrete NanoLuc (Nluc) in the mouse gut. The molecular toolkit we present here should provide an accessible framework for generating living therapeutic and diagnostic machines from highly relevant human commensal *Bacteroides* species.

## Results

### P_BfP1E6_-RBS8 promoter/RBS drives strong and reproducible protein secretion in *B. theta*

To establish a set of secretion tools for members of the *Bacteroides* genus, we selected *B. theta* as the starting point based on its prevalence and abundance in the human gut as well as the large body of knowledge surrounding this species, including a substantial collection of well characterized genetic tools^16–24^. To ensure robust and reproducible results, we first sought to establish a framework for evaluation of protein expression and secretion across diverse samples. We identified a core set of three native *B. theta* proteins previously shown to be highly secreted, each with a different N-terminal signal sequence: BT_2472 (Sec/SPI SP), BT_3382 (lipoprotein SP), and BT_3769 (no SP identified; secretion mechanism unknown)^43^. To identify optimal genetic parts for reproducible and detectable protein secretion, we tested each protein using three different promoter/ribosome-binding site (RBS) pairs **(Fig. 1A)**. For strong, constitutive expression, we used a previously characterized *Bacteroides fragilis* phage promoter paired with either its original ribosome-binding site (RBS_phage_) or a modified version with the highest observed activity amongst reported variants (RBS8)^20^. For more precisely controlled protein expression, we used a tightly regulated anhydrotetracycline (aTc)-inducible promoter (P1T_DP_-GH023)^18^. To evaluate the performance of the different constructs and to determine the best timepoint for measuring extracellular protein accumulation in future studies, we expressed each protein from each expression plasmid and monitored bacterial growth and secretion in *B. theta* liquid culture for 48 hours **(Fig. 1B)**. Despite similar growth kinetics between samples, we only observed high levels of secretion from BT_3382, and only with the strong, constitutive phage promoter/RBS pairs. BT_3769 only achieved levels of secretion above background when expression was driven by P_BfP1E6_-RBS8, however the signal was more than two-fold lower than that measured for BT_3382 with the same promoter/RBS. Interestingly, P1T_DP_-GH023 did not drive high enough protein expression to result in detectable levels of secretion for any of the three proteins, even after increasing the concentration of aTc from 100 ng/mL to 200 ng/mL **(Fig. 1B)**. Similarly, no expression construct produced detectable levels of secreted BT_2472 **(Fig. 1B)**. While the cause is unclear, it is possible that the C-terminal domain of this protein is involved in secretion and was compromised by the 3xFLAG tag that was included for immunodetection.

**Figure 1.**
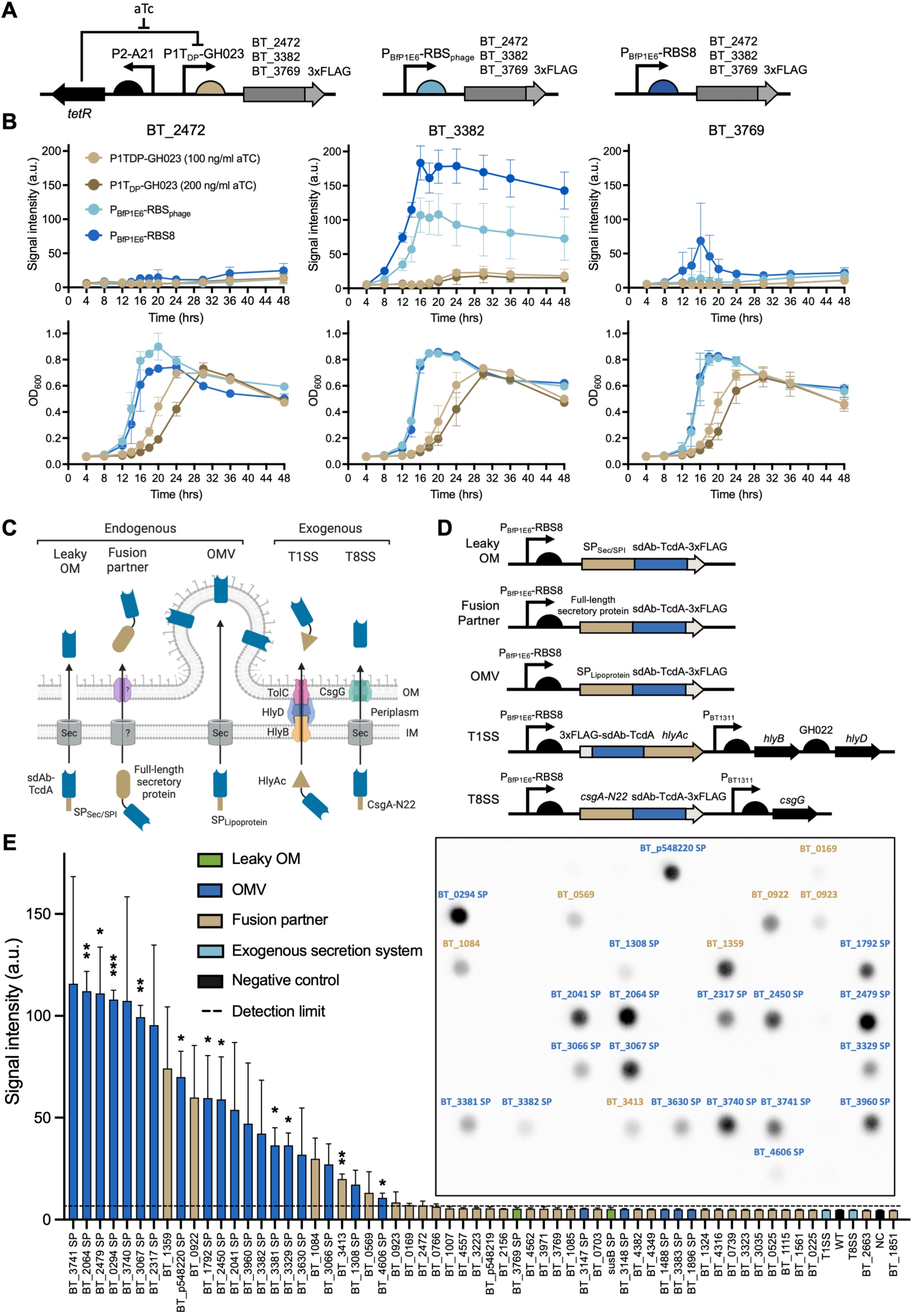
Engineering *B. theta* to secrete sdAb-TcdA. (A) Design of genetic constructs for protein expression and secretion. (B) Protein secretion (top row) and growth (bottom row) of *B. theta* expressing BT_2472, BT_3382, and BT_3769 with their native secretion signals, measured from supernatant samples taken every 4 hrs for 48 hrs. Protein levels were measured by dot blot and bacterial growth was measured by optical density at 600 nm (OD_600_). For strains using the P1T_DP_-GH023 promoter, the inducer aTc (100 or 200 ng/ml) was included in the medium at the time of inoculation. Error bars represent standard deviation of three biological replicates. a.u., arbitrary units. (C) Schematic representation of secretion strategies explored in this study. (D) Design of genetic constructs for secretion carrier screening in *B. theta*. (E) Relative levels of secretion of sdAb-TcdA in culture supernatants of *B. theta* harboring sixty different expression/secretion constructs, measured by dot blot. Inset shows representative dot blot with effective secretion carriers (above detection limit) labeled. Detection limit (dotted line) was set at the signal intensity of the faintest dot visible by unaided eye on the membrane, which is around 7 arbitrary units (a.u.). Error bars represent one standard deviation of triplicate biological samples. Significance was determined using unpaired two-tailed Welch’s *t* test. ∗p < 0.05, ∗∗p < 0.01, ***p < 0.001. WT, wild-type *B. theta*; NC, negative control *B. theta* expressing sdAb-TcdA with no secretion carrier fusion.

For both BT_3382 and BT_3769, we observed peak extracellular protein accumulation when *B. theta* cultures grew to late log phase (16-20 hr, OD_600_ 0.6-0.8). Beyond this point however, the amount of BT_3769 in the culture media rapidly dropped to undetectable levels within 8 hours, whereas the level of BT_3382 only dropped by ∼15% when measured 48 hr later **(Fig. 1B)**. We suspect that the persistently high levels of BT_3382 are due to higher protein stability rather than continued production and secretion by *B. theta* during stationery growth as BT_3382 is reported to be enriched in OMVs^43^, which may lead to higher thermostability^44,45^.

Based on these results, we decided to 1) use P_BfP1E6_-RBS8 as our baseline promoter/RBS pair to drive expression of all constructs moving forward since it resulted in the highest observed secretion levels in this preliminary screen, and 2) collect all supernatant samples between late log and stationary phase of growth to ensure consistent detection of secreted products within the predicted window of protein stability for all samples.

### Identification of secretion carrier candidates from *B. theta* endogenous machinery and *E. coli* exogenous machinery

To continue developing secretion toolkit, we next sought to identify signal peptides, full-length proteins, or protein domains to serve as “secretion carriers” that promote extracellular export of heterologous proteins from *B. theta*. Typical approaches either utilize endogenous secretion machinery with homology to known systems or introduce exogenous secretion systems/tags from other bacterial strains^42^. Because most previously characterized secretion systems in Gram-negative bacteria are either incomplete or not conserved in the *B. theta* genome^37,38^, the endogenous secretion systems of *B. theta* are still poorly understood. To circumvent this limitation, we identified three secretion strategies that are typically applicable to all Gram-negative bacteria – leaky outer membrane (OM), fusion partner, and outer membrane vesicle (OMV) (**Fig. 1C**) – and searched for endogenous secretion carrier candidates within each of these categories in *B. theta* genome. To ensure thorough investigation of all potential solutions for promoting secretion of heterologous proteins in *B. theta*, we also selected two highly studied exogenous secretion systems from *Escherichia coli* (*E. coli*) -T1SS and T8SS **(Fig. 1C)** and transferred them into *B. theta* for testing as well. To evaluate the efficiency of the secretion carriers, we selected a single candidate protein to serve as our standard secretion cargo. Because our goal is to establish a toolbox to enable the development and implementation of *Bacteroides* species as living therapeutics in their natural gut environment, we selected a clinically relevant single domain antibody (sdAb) that targets Toxin A (TcdA) from *Clostridioides difficile*^46^, a prominent and challenging gastrointestinal pathogen^47^. Compared to full-length antibodies, the small size and structural simplicity of sdAbs allows them to be more easily expressed by bacteria and results in higher thermal and proteolytic stability in the harsh gut environment.

The leaky OM strategy relies on transport of proteins into the periplasm via Sec pathway, followed by secretion to the extracellular space through natural OM leakage^42^ **(Fig. 1C)**. We selected two Sec/SPI SPs from two *B. theta* endogenous proteins as secretion carriers for leaky OM strategy: SusB, a periplasmic protein of the well-studied *B. theta* starch utilization system (Sus)^48^ with a Sec/SPI SP, and BT_3769, which also has a Sec/SPI SP and was previously identified as highly secreted^43^. The fusion partner strategy requires heterologous cargoes to be genetically fused with full-length native secretory proteins for co-transportation out of the cell^42^ **(Fig. 1C)**, without the need to understand the underlying secretion mechanisms. To search for secretion carriers for fusion partner strategy, we referred to a previous study characterizing endogenous protein abundance in different fractions (inner membrane (IM), OM, OMV pellets (OMVp), and OMV-free supernatants (SUP)) of *B. theta* liquid culture^43^ and identified 33 candidates with high abundance in the OMVp, SUP, or both fractions **(Table S1, S2 and Methods)**. Finally, the OMV strategy is based on the recent discovery of the lipoprotein export signal (LES), a five-residue conserved motif that immediately follows the lipoprotein SP cleavage site (+2 to +6) in many native OMV-enriched lipoproteins in *B. theta*^43,49,50^. We hypothesized that adding a lipoprotein SP with an identifiable LES to heterologous cargoes would enable their secretion via OMVs **(Fig. 1C)**. Using a similar approach as fusion partner strategy, we identified the SPs of 24 lipoproteins enriched in either the OMVp or the combined (OMVp+SUP) fractions of *B. theta* liquid culture as the secretion carriers for OMV strategy^43^ **(Tables S1, S2 & Methods)**. For the above identified endogenous *B. theta* secretion carriers (candidate SP or full-length carrier protein), we fused them to the N-terminus of sdAb-TcdA and included a C-terminal 3xFLAG tag for detection **(Fig. 1D).**

For the strategy of introducing exogenous secretion machinery, we selected two systems with a small genetic size, few components, and simple regulation to ensure the highest chance of success: the hemolysin system (T1SS) of uropathogenic *E. coli* (UPEC)^51^ and the curli system (T8SS) from *E. coli* K-12^52^, which have both been used successfully for heterologous protein secretion in non-native hosts^53,54^. The hemolysin system contains HlyB, HlyD, and TolC, which form the secretion channel, and HlyA, which is the cognate secreted product used to drive co-transport of protein cargoes via fusion to its C-terminal domain (HlyAc) **(Fig. 1C, Table S2).** We fused HlyAc domain at the C-terminus of the sdAb-TcdA and inserted the HlyB and HlyD genes into the same plasmid in a polycistronic format under the control of a strong native *B. theta* constitutive promoter P_BT1311_ and *B. theta* GH022 RBS^19^ in a polycistronic format **(Fig. 1D).** We did not include TolC in this construct because it is conserved in most bacterial species^55^ and several putative homologs exist in *B. theta*^56^. The curli secretion system (T8SS) is even simpler than the hemolysin system, requiring only a CsgG transport protein (expressed by P_BT1311_) and an N-terminal fusion of the first 22 amino acids of the cognate secreted product (CsgA-N22) to the cargo protein ^54^ **(Fig. 1C, 1D, Table S2).**

### Native *B. theta* secretion carriers enable high-level extracellular export of sdAb-TcdA

Of the sixty-one secretion carriers we identified **(Table S2)** and fused to the sdAb-TcdA, all constructs were successfully cloned and conjugated into *B. theta* except for BT_3434, which appeared to be lethal in *E. coli* DH5α. To determine the secretion efficiency of each of the other sixty secretion carriers, we grew *B. theta* transconjugant cultures to late-log phase and measured the abundance of sdAb-TcdA in culture supernatant by dot blot **(Fig. 1E).** Twenty-six (43%) of the sixty secretion carriers produced visible signal and were thus considered “effective.” However, no candidate from the leaky OM nor the exogenous secretion system approaches were represented in this group. These results are not entirely unexpected, as the leaky OM approach usually requires membrane-disrupting methods for full efficacy^57,58^, and the *E. coli* T1SS and T8SS may require unknown accessory proteins or regulators^52,59,60^ that are not conserved in *B. theta*. Of the twenty-six effective secretion carriers, seven (27% of effective candidates, 12% of total) are full-length fusion partner proteins (six with Sec/SPI SPs and one with a lipoprotein SP) and nineteen (73% of effective candidates, 32% of total) are lipoprotein SPs. The seven effective fusion partners represent only 22% of the thirty-two total fusion partners tested. The relatively low success rate of this class of secretion carrier could be improved with optimization of the fusion, e.g., truncation mutants, alternate orientations (N-, C-, or in-frame internal fusions), etc.^61–63^. Conversely, the efficient secretion of sdAb-TcdA observed for 79% of the lipoprotein SPs (19/24) supports our hypothesis that lipoprotein SPs with LES sequences may be able to drive OMV-mediated secretion of heterologous proteins.

### A positively charged region and a length-restricted hydrophobic region are critical for effective heterologous protein secretion by lipoprotein SPs

Most of the effective secretion carriers we identified were lipoprotein SPs of OMV secretion strategy, however, five *B. theta* lipoprotein SPs that we tested did not effectively mediate secretion of sdAb-TcdA: BT_1488, BT_1896, BT_3147, BT_3148, and BT_3383 **(Fig. 1E).** To determine if effective lipoprotein SPs harbor other unique features in addition to the LES, we further analyzed the amino acid sequences of our lipoprotein SP collection. These SPs are composed of a positively charged N-terminal region (n), a central hydrophobic region (h), a cysteine residue after the cleavage site (+1), and a negatively charged LES motif (+2 to +6). Interestingly, we found that the backbones (n- and h- regions) of the five ineffective lipoprotein SPs are either very short (∼10 residues) or very long (∼40 residues), compared to the backbones of effective lipoprotein SPs (16-34 residues) **(Fig. 2A).** All five ineffective lipoprotein SPs lack a positively charged n-region and BT_3383 SP also has no clear h-region **(Fig. 2B).** These results suggest that, in addition to the LES, the presence of positively charged residues in the n-region, as well as a minimum length of the h-region may be critical factors that drive lipoprotein SP-mediated secretion in *B. theta*.

**Figure 2.**
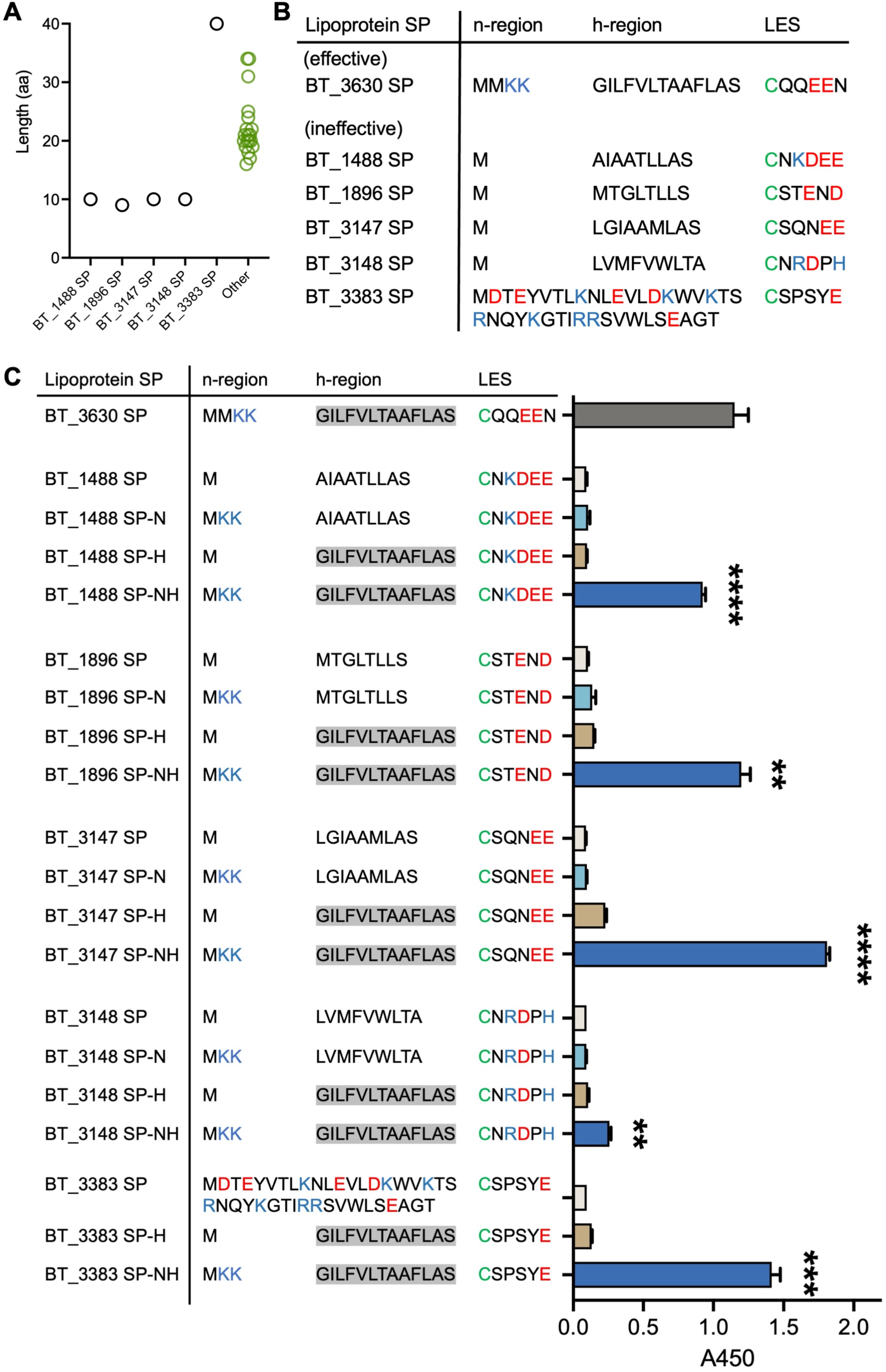
Rational engineering enables ineffective lipoprotein SPs to secrete sdAb-TcdA. (A) Comparison of the length of five ineffective (black) and nineteen effective (green) lipoprotein SP sequences. (B) Comparison of amino acid sequences of the five ineffective lipoprotein SPs with a prototypical effective sequence. Positively charged residues are colored blue, negatively charged residues are colored red, and cysteine residue cleavage sites are colored green. (C) Secretion of sdAb-TcdA driven by native (SP) and domain-swapped (SP-N, SP-H, and SP-NH) ineffective lipoprotein SPs, measured by ELISA. BT_3630 SP-sdAb-TcdA is included as positive control. Residue coloring is the same as for (B), and hydrophobic regions are boxed in gray. Error bars represent one standard deviation of triplicate biological samples. Significance was determined using unpaired two-tailed Welch’s *t* test. ∗∗p < 0.01, ***p < 0.001, ****p < 0.0001

To test this hypothesis, we swapped the n- and h-regions of the five ineffective SPs with those from an effective SP to see if we could improve their secretion efficiency through rational design. We chose the SP from BT_3630 as our standard based on its layout of charged and hydrophobic regions, which is broadly representative of the collection of lipoprotein SPs that we identified as effective **(Fig. 2B, 2C).** For each of the five ineffective lipoprotein SP sequences, we made the following three SP variants and fused them to the sdAb-TcdA: 1) added two N-terminal lysines immediately after the start codon to introduce the positively charged region (SP-N), 2) replaced the hydrophobic region with the one from BT_3630 SP (SP-H), or 3) introduced both modifications (SP-NH) **(Fig. 2C).** The only variant that was not generated was the SP-N version of the BT_3383 SP; because it does not have an obvious h-region we concluded that addition of an n-region would not be sufficient to improve its secretion capability.

We observed enhanced secretion of the sdAb-TcdA only for SPs that had both an added N-terminal charged domain and a swapped hydrophobic region (SP-NH variants) **(Fig. 2C),** suggesting that both regions are necessary, but neither is sufficient to drive high-level secretion in *B. theta*. Notably, the increase in sdAb secretion measured for the BT_3148 SP-NH variant was substantially lower than for the other SP-NH variants **(Fig. 2C).** Inspection of the LES sequence revealed positively charged amino acids in the +3 and +6 positions, which is consistent with previous findings showing that interspersed positively charged amino acids can offset LES- mediated OMV secretion ^43^. These results begin to define the requirements for the presence and placement of charged and hydrophobic residues in *B. theta* lipoprotein SP sequences that can derive heterologous protein secretion.

### *B. theta* secretion carriers mediate export of multiple types of functional protein cargoes

Toward our goal of establishing a flexible toolbox to enable efficient secretion of diverse heterologous protein cargoes, we next tested the ability of the twenty-six effective secretion carriers **(Fig. 1E)** to secrete six additional proteins with therapeutic and/or diagnostic functions. Three of the six proteins are disease-targeting antibody fragments, including an sdAb targeting tumor necrosis factor alpha (sdAb-TNFα)^64^, an antigen associated with chronic conditions such as inflammatory bowel disease (IBD)^65^; another sdAb targeting epidermal growth factor receptor (sdAb-EGFR)^66^, commonly overexpressed in colon cancer^67^; and a single-chain variable fragment targeting human epidermal growth factor receptor-2 (scFv-HER2)^68^, mainly known for its role in breast cancer, but also implicated in colon cancer^69^. This set of proteins allows us to evaluate secretion efficiency across diverse antibody fragment formats while still focusing on targets relevant to gastrointestinal delivery by engineered living therapeutics. We selected the other three proteins to provide a diverse set of reporter functions: NanoLuc (Nluc)^70^, enhanced green fluorescent protein (EGFP)^71^, and β-lactamase (BLac)^72^, which yield quantifiable outputs of luminescence, fluorescence, or colorimetric signal, respectively.

Each of these six cargo proteins were fused to each of the twenty-six secretion carriers, resulting in one hundred and fifty-six new carrier-cargo pairs. With the exception of EGFP, all cargoes were secreted from *B. theta* and accumulated at varying levels in culture supernatants via different secretion carriers **(Fig. 3A).** We suspect that the exceptionally low levels of EGFP detected in culture media may be due to rapid folding of this protein, which has been reported to stall the translocon protein during secretion process^73^. For the other cargoes, we observed considerable variability in secretion efficiency both between different secretion carriers as well as amongst cargoes secreted by the same carrier. In contrast to EGFP, we detected secreted Nluc at high levels across the majority of secretion carriers, suggesting that Nluc can be broadly used as a highly sensitive reporter for measuring secretion efficiency. Compared to the other proteins tested, Nluc has a higher solubility and a more acidic isoelectric point (**Table S3**), both of which have been reported to enhance protein secretion^74,75^ and hence might account for its high-level secretion. Interestingly, although all sdAbs share similar structural framework^76^, the three sdAbs tested here were not secreted at consistent levels by the same secretion carriers. It has been previously reported that cargo-specific interactions with signal peptides can indirectly impact secretion by influencing other cellular processes such as protein biosynthesis, folding kinetics, and structural stability^77^, which could explain some of the variability that we observed.

**Figure 3.**
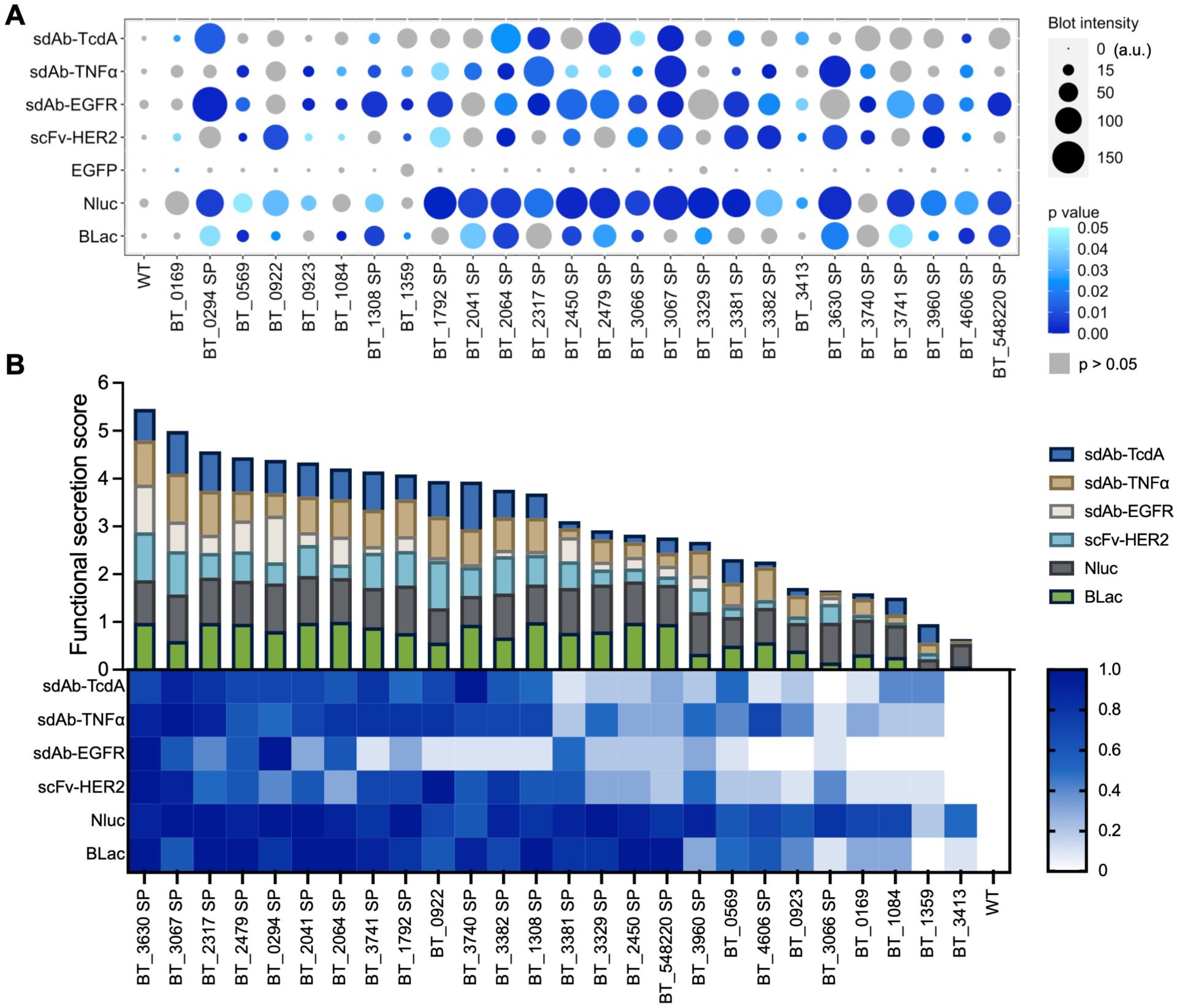
*B. theta*-derived secretion carriers function across multiple heterologous proteins. (A) Relative levels of antibody fragments and reporter proteins secreted into culture supernatant by *B. theta* secretion carriers. Bubble size corresponds to average blot intensity of triplicate experiments with p < 0.05 indicated by the blue color scale and p > 0.05 shown in gray. Significance was determined using unpaired two-tailed Welch’s *t* test. (B) Functional assays of antibody fragments and reporter proteins secreted into culture supernatant by *B. theta* secretion carriers. Binding of antibody fragments (sdAbs and scFv) to their respective targets was determined by ELISA. Enzymatic activity of reporter proteins Nluc and BLac was determined by bioluminescence assay and colorimetric assay, respectively. Following log transformation of luminescence data, all functional assay readouts were converted to values between zero and one by cargo-wise min-max normalization.

Finally, to verify that the secreted protein products were properly folded and not otherwise functionally disrupted by fusion to the secretion carriers, we performed functional assays to measure the antigen binding or enzymatic activity of each of the secreted cargo proteins in *B. theta* culture supernatants. Because the readouts of these functional assays are not equivalent across cargoes **(Fig. S1),** we scaled the output values to a normalized range of zero to one, corresponding to the lowest and highest readout for each cargo across the twenty-six secretion carriers **(Fig. 3B**, lower panel). We then created a “functional secretion score” for each carrier by summing the normalized functional outputs for all six cargo proteins for each secretion carrier **(Fig. 3B**, upper panel). These results revealed several lipoprotein SPs that are broadly effective for secreting diverse cargoes with BT_3630 SP and BT_3067 SP emerging as the most consistent and robust broadly active secretion carriers.

### *B. theta*-derived secretion carriers function across multiple *Bacteroides* species

Toward the goal of developing universal secretion tools for the *Bacteroides* genus, we next sought to evaluate the *B. theta*-derived secretion carriers in other *Bacteroides* species. We selected the ten carriers with the highest functional secretion scores **(Fig. 3B)** and measured the level of secretion **(Fig. 4A)** and functionality **(Fig. 4B)** of each of the six cargo proteins when expressed in three different *Bacteroides* species: *B. fragilis, B. ovatus,* and *B. vulgatus.* We were unable to generate *B. fragilis* transconjugants for six of the carrier-cargo pairs, which may be due to low conjugation efficiency in this species^20^ or lethal intracellular aggregation of protein cargoes^78^. For all other *Bacteroides* transconjugants, the results mirrored those observed in *B. theta* **(Fig. 3)**; secretion efficiency varies not only between cargoes but also between species. Among the *Bacteroides* species tested, *B. ovatus* generally demonstrated the highest secretion levels for any given carrier-cargo pair, which may be due to the closer phylogenetic relationship between *B. theta* and *B. ovatus*^18^. As we observed with *B. theta*, Nluc generally had the highest secretion levels among the six cargo proteins across all three *Bacteroides* species. In contrast, efficient secretion of sdAb-EGFR and scFv-HER2 appears to be restricted to only a few selected secretion carriers.

**Figure 4.**
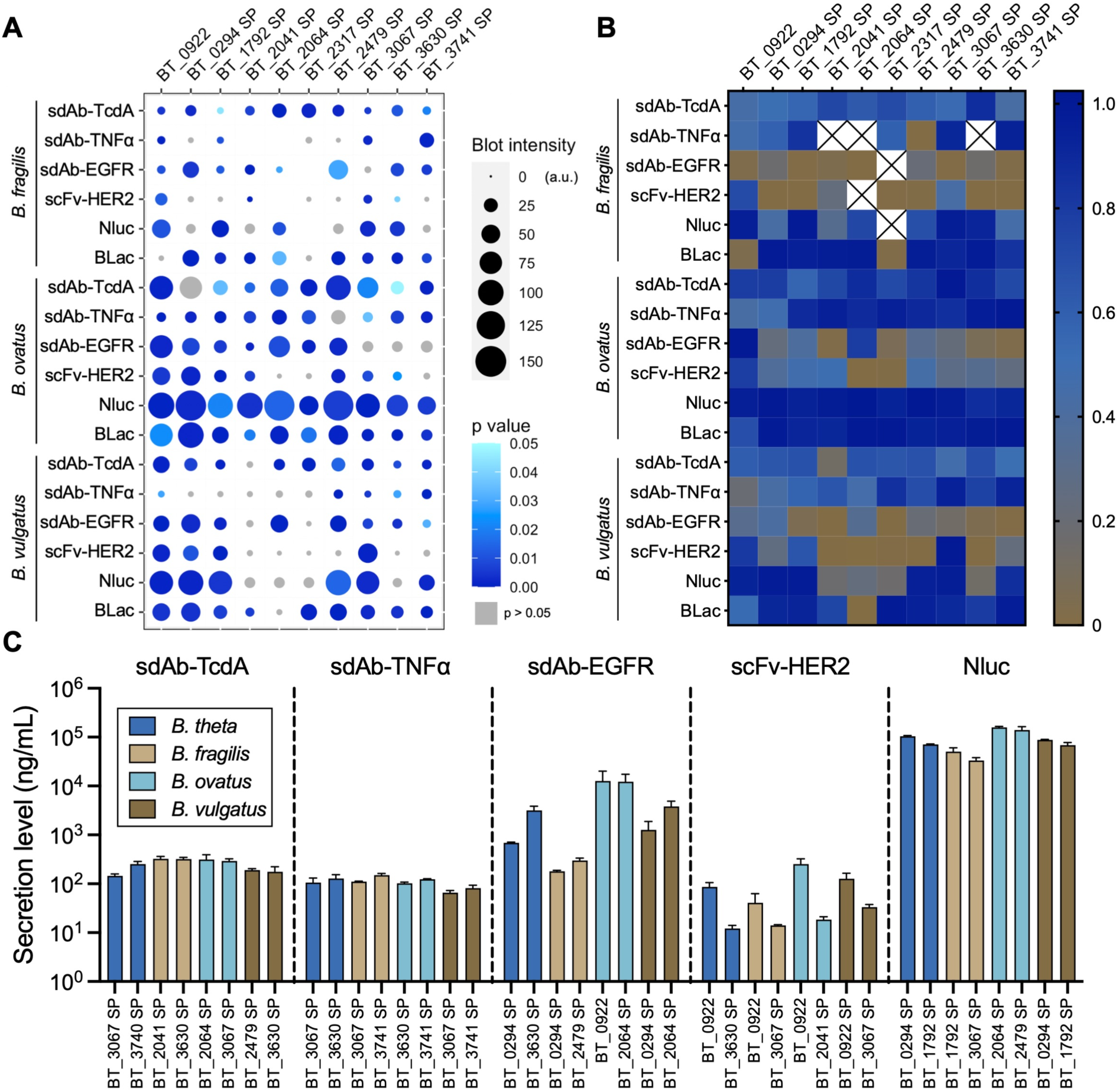
*B. theta*-derived secretion carriers mediate export of diverse, functional cargoes from multiple *Bacteroides* species. (A) Relative levels of six cargo proteins detected in the culture supernatants of three *Bacteroides* species, driven by each of the ten highest performing native *B. theta* secretion carriers. Bubble size corresponds to average blot intensity of triplicate experiments with p < 0.05 indicated by the blue color scale and p > 0.05 shown in gray. Significance was determined using unpaired two-tailed Welch’s *t* test. (B) Functional assays of antibody fragments and reporter proteins secreted into culture supernatant by secretion carriers. Binding of antibody fragments (sdAb-TcdA, sdAb- TNFα, sdAb-EGFR, and scFv-HER2) to their respective targets was determined by ELISA. Enzymatic activity of reporter proteins (Nluc and BLac) was determined by bioluminescence assay and colorimetric assay, respectively. Following log transformation of luminescence data, all functional assay readouts were converted to values between zero and one by cargo-wise min- max normalization. All functional assays were performed in triplicate. (C) Quantification of protein secretion titers mediated by the two secretion carriers that yielded the highest functional protein levels of each cargo in each species. Error bars represent the standard deviation of triplicate experiments.

After validating the heterologous protein secretion capability of the *B. theta*-derived secretion carriers across four *Bacteroides* species, we next sought to quantify secretion titers in these species. For these measurements, we selected five of our seven cargo proteins: four antibody fragments and Nluc. As noted above, EGFP was not secreted **(Fig. 3A)** and, while BLac is highly secreted, it reaches saturation in the enzymatic assay too quickly to be suitable for quantification **(Fig. 3B, 4B, S1)**, thus we excluded these two reporters from this assessment. For each of the other five cargoes, we identified the two secretion carriers that yielded the highest functional protein levels in culture supernatant of each cargo in each species **(Fig. 4B)** and selected these carrier-cargo-species combinations for our analysis. Cultures were grown to late-log phase and the level of protein in each culture supernatant was quantified by comparison to standard curves of known concentrations of purified proteins **(Fig. 4C).** For the subset of secretion carriers tested, we observed secretion titers within a relatively narrow range across all four *Bacteroides* species for sdAb-TcdA (145-320 ng/mL) and sdAb-TNFα (65-150 ng/mL). Conversely, the secretion levels of sdAb-EGFR (180-12000 ng/mL), scFv-HER2 (10-250 ng/mL), and Nluc (33000-158000 ng/mL) were much more variable between secretion carriers, *Bacteroides* species, or both.

### Modified inducible expression system yields enhanced protein secretion

Having successfully established an approach to enable heterologous protein secretion from *B. theta* and other *Bacteroides* species, we next sought to engineer additional layers of flexibility, control, and enhancement using an inducible gene expression system. In our initial studies, we observed that the aTc-inducible P2-A21-tetR-P1T_DP_-GH023 expression cassette resulted in much lower secretion than P_BfP1E6_-RBS8 **(Fig. 1B),** presumably due to lower expression. To generate an inducible system capable of achieving much higher expression levels, and thus much higher secretion levels, we introduced two modifications aimed at enhancing activity. First, we replaced the GH023 RBS with the A21 RBS **(Fig. 5A),** which was previously identified as the strongest RBS tested amongst a collection of *Bacteroides* RBS sequences when paired with the P1T_DP_ promoter^18^. Because the original construct already contained an A21 RBS sequence **(Fig. 1A),** we replaced the TetR-driving P2-A21 promoter/RBS with P_BT1311_ and its native RBS **(Fig. 5A)** to avoid issues such as unwanted homologous recombination events between identical RBS sequences^79^.

**Figure 5.**
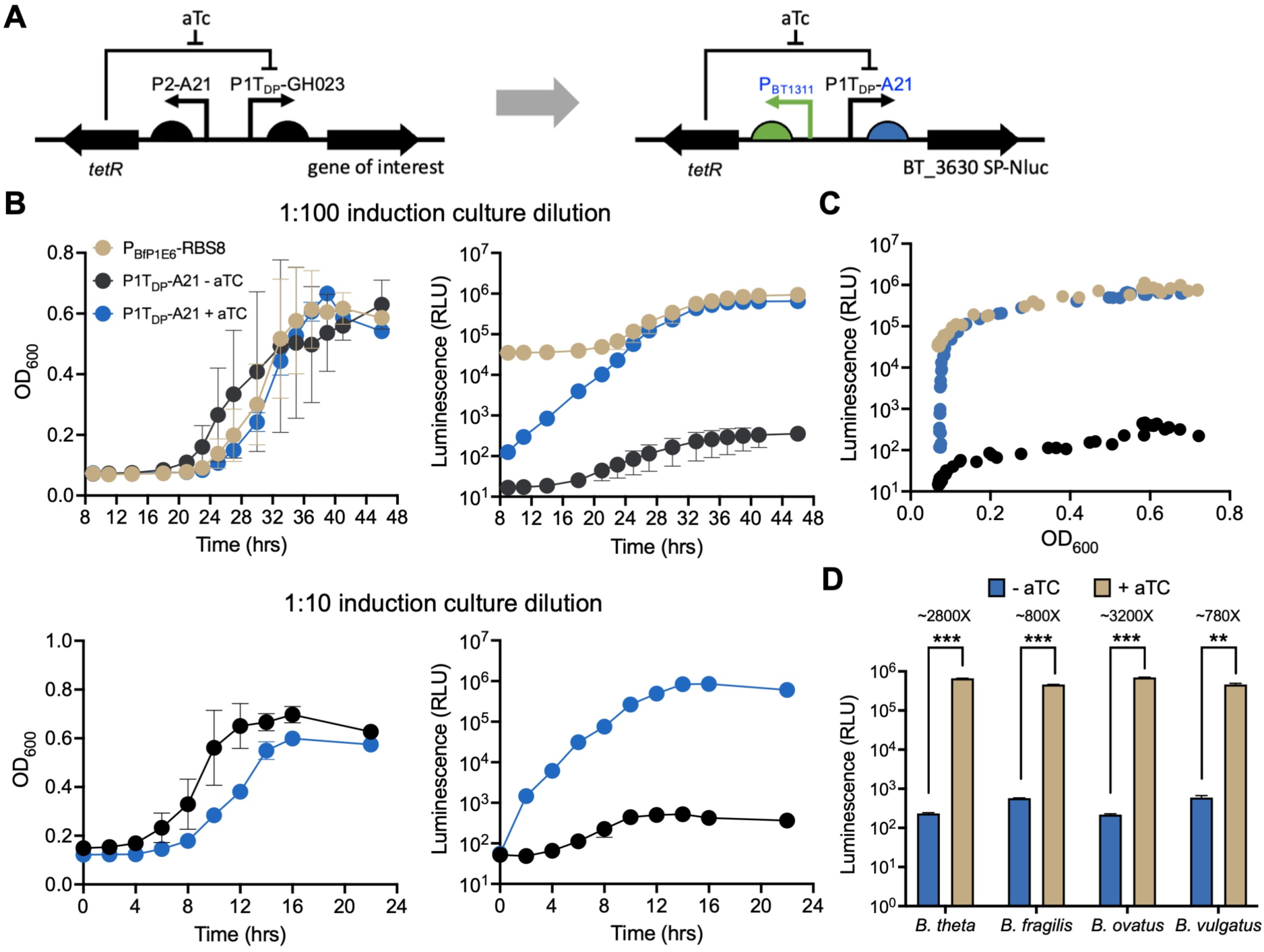
Development of a strong, aTc-inducible expression cassette for enhanced control of protein secretion across multiple *Bacteroides* species. (A) Low-activity promoter and RBS sequences in the original P2-A21-tetR-P1T_DP_-GH023 inducible expression cassette (left)^18^ were replaced with high-activity variants to generate the modified P_BT1311_-tetR-P1T_DP_-A21 expression cassette (right). (B) Modified inducible expression cassette drives expression of Nluc reporter at levels similar to high-level constitutive promoter P_BfP1E6_-RBS8 in cultures diluted at 1:100 (top) or 1:10 (bottom). (C) Correlation between bacterial growth and Nluc secretion levels shown in Fig. 5B, top. (D) Modified inducible system mediates tightly controlled protein expression across multiple *Bacteroides* species. For all experiments, luminescence measurements were obtained from clarified culture supernatant and thus represent the secreted fraction of total expressed Nluc. Nluc secretion in this study was mediated by the highly active secretion carrier BT_3630 SP. Error bars represent one standard deviation of triplicate biological samples. Significance was determined using unpaired two-tailed Welch’s *t* test. ∗∗p < 0.01, ***p < 0.001, ****p < 0.0001

To measure the activity of our enhanced inducible expression system, we fused Nluc with the high-efficiency secretion carrier BT_3630 SP and generated two expression/secretion constructs: one driven by the aTc-inducible P1T_DP_-A21 promoter, and one driven by the high-activity constitutive P_BfP1E6_-RBS8 as both a positive control and reference point for high-level expression **(Fig. 5B, top)**. To quantify secretion, we induced freshly diluted (1:100) overnight cultures with 100 ng/mL aTc and performed bioluminescence measurements from 8 to 46 hr post-induction. Induction of Nluc expression from the enhanced P1T_DP_-A21 promoter resulted in secretion levels up to 2000-fold higher than uninduced controls and ∼70% of the P_BfP1E6_-RBS8 constitutively expressed control, achieving the same order of magnitude of secretion level as the strongest promoter/RBS pair, P_BfP1E6_-RBS8. **(Fig. 5B, top)**. Because the *B. theta* cultures demonstrated a period of initial slow growth following a 1:100 dilution of the overnight cultures into fresh induction medium, we repeated the experiment using cultures diluted at 1:10. These samples achieved late-log phase growth after only 14 hr, compared to nearly 35 hr for the samples diluted at 1:100, but reached nearly the same maximal secretion levels **(Fig. 5B, bottom)**, which suggests that the density of the culture does not affect the induction of this promoter. As we observed in our initial growth experiments (**Fig. 1B**), secretion levels are highly correlated with the growth phase of liquid cultures (**Fig. 5C**). Furthermore, the growth curves for bacteria harboring the inducible P1T_DP_-A21 expression cassette are similar in both the presence and absence of the inducer aTc **(Fig. 5B),** suggesting that the inducible system does not impart an obvious metabolic burden on *B. theta*. To verify the portability of this modified promoter, we repeated the experiments in all four different *Bacteroides* species and observed that the enhanced inducible expression system yields similar levels of secreted Nluc, and similar fold-induction levels compared to uninduced samples, across all four species tested **(Fig. 5D).** Interestingly, for *B. fragilis* and *B. vulgatus*, this promoter gave rise to slightly lower secretion levels in induced cultures and slightly higher apparent expression leakage in uninduced cultures, resulting in overall lower fold-induction levels for these two species. This difference, similar to what we observed for overall secretion carrier performance across *Bacteroides* species **(Fig. 4),** may also be linked to their evolutionary distance from the other two species.

### Different secretion carriers mediate distinct post-secretion extracellular fate of protein cargoes

Toward our goal of reproducibly delivering therapeutic proteins into specific physiological niches such as the gut lumen, we next sought to investigate the post-secretion extracellular fate of heterologous proteins exported using our platform. Because we expect OMV-associated proteins to have fundamentally different characteristics than freely soluble proteins, such as thermostability, protease resistance, bioavailability, and dissemination to other body sites^80^, precise determination of the extracellular destination mediated by different secretion carriers is required to fully understand and optimize our platform. Based on the high secretion levels and high sensitivity observed in earlier experiments, we selected Nluc as the secretion cargo for these studies. From our collection of twenty-six effective secretion carriers **(Fig. 1E),** we selected four candidates with diverse structures and Nluc secretion efficiency **(Fig. 3B, S1)** across fusion partner and OMV strategies for further investigation: BT_0169 and BT_0569 (fusion partner strategy; full-length secretion carriers with Sec/SPI SPs); BT_0922 (fusion partner strategy; full-length secretion carriers with lipoprotein SP); BT_3630 SP (OMV strategy; lipoprotein SP).

To determine the extracellular fate of Nluc when secreted by these four carriers, we grew late-log phase liquid cultures of *B. theta* expressing each carrier-Nluc fusion, separated the cell pellets (P) from the total supernatants (T), then further separated the total supernatants into the soluble (S) and insoluble OMV (O) fractions. Cell pellets were concentrated 2.5-fold and OMV fractions were concentrated 20-fold during the extraction process. We measured the Nluc protein abundance **(Fig. 6A)** and luminescence activity **(Fig. 6B)** in each fraction. Consistent with the differences in carrier-specific Nluc secretion efficiency observed in earlier experiments **(Fig. 3)**, western blot analysis revealed that the majority of the BT_0169-Nluc and BT_0569-Nluc was retained in the cell pellets, whereas BT_0922-Nluc and BT_3630 SP-Nluc were mostly secreted, although with different abundances in different fractions. BT_0922-Nluc appeared to have been cleaved, showing a faint band at its expected molecular weight of ∼60 kDa in the cell pellet as well as OMV fractions while only a smaller, ∼23 kDa product was detected in all other culture fractions. In the luciferase assay we found that, after separating out the OMVs, the soluble fraction of BT_0169-Nluc accounted for ∼80% of the luminescence observed in the total supernatant **(Fig. 6B),** suggesting that BT_0169-Nluc is mainly secreted in a freely soluble form. The luminescence signals of the OMV-free soluble fractions of BT_0569-Nluc, BT_0922-Nluc, and BT_3630 SP-Nluc account for ∼50% of the total supernatant signal, suggesting that these three secretion carriers mediate extracellular export of freely soluble and OMV-bound cargoes in roughly equal proportions. However, the luminescence signal from the concentrated OMV fractions was substantially higher for BT_0569 than for the other three secretion carriers. Remarkably, BT_3630 SP-Nluc appeared in both soluble and OMV fractions, suggesting that lipoprotein SPs secrete heterologous proteins through not only OMV-dependent but also OMV-independent pathways, which has been reported previously^81–83^, but, to our knowledge, never for lipoprotein SPs nor in *Bacteroides* species.

**Figure 6.**
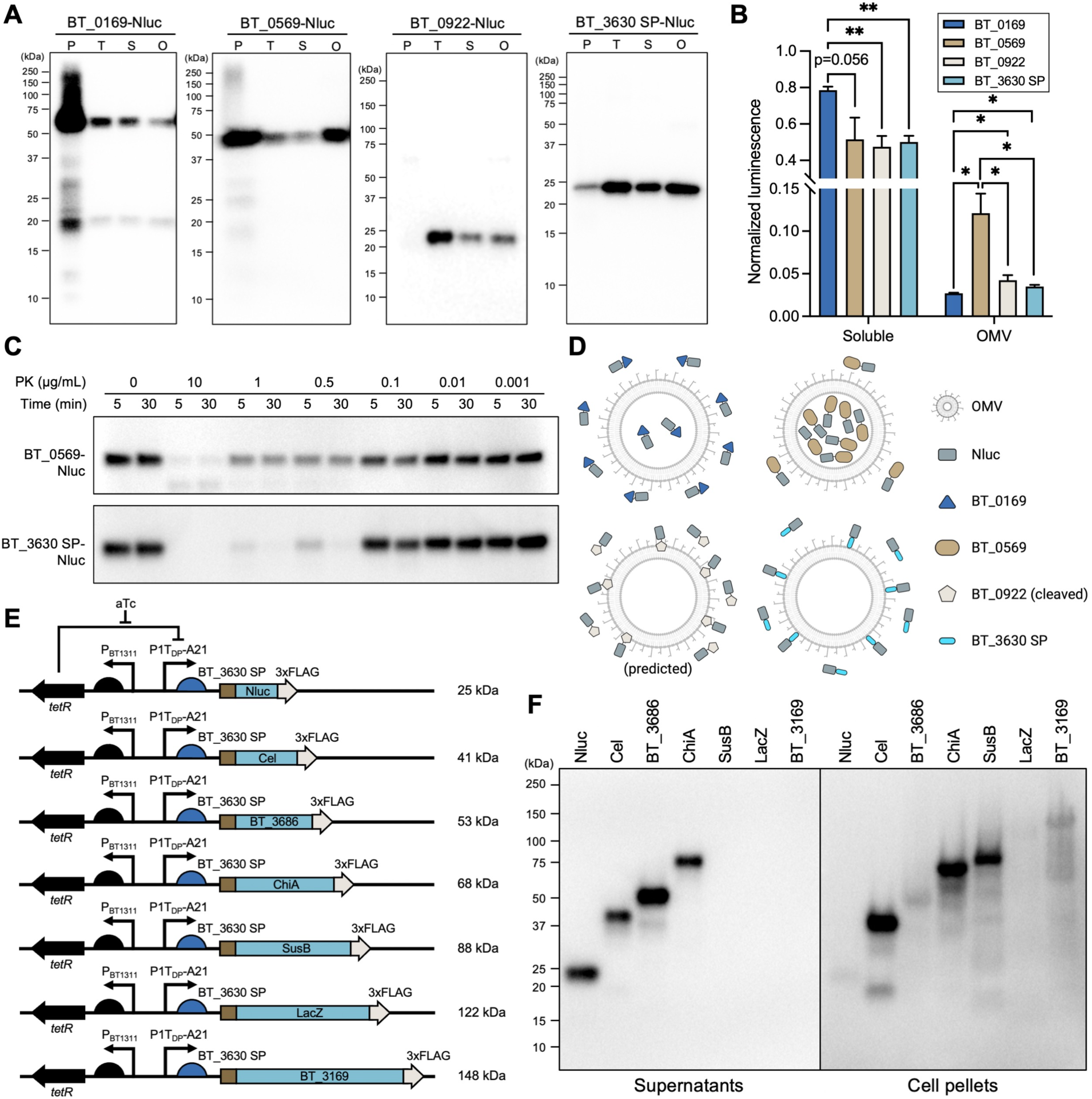
Characterization of the post-secretion extracellular fate and size limit of cargo proteins for secretion carriers. (A) Western blot analysis of Nluc abundance in different fractions of *B. theta* liquid cultures expressing four carrier-Nluc constructs. P, cell pellet; T, total supernatant; S, soluble fraction of total supernatant; and O, OMV fraction of total supernatant. P and O are 2.5-fold and 20-fold more concentrated than supernatant of equivalent volume, respectively. (B) Enzymatic activity of secreted Nluc in soluble and OMV fractions, measured by luminescence assay. To normalize the efficiency difference between the four secretion carriers, the luminescence in soluble and OMV fractions was divided by the luminescence in total supernatants to calculate the relative abundance of secreted Nluc in soluble and OMV fractions. Normalized luminescence of concentrated OMV fractions was divided by 20 to correct for concentration during sample prep. Error bars represent one standard deviation of triplicate biological samples. Significance was determined using unpaired two-tailed Welch’s *t* test. ∗p < 0.05, **p < 0.01 (C) Western blot analysis of proteinase K assay of OMV fractions from *B. theta* cultures expressing BT_0569- Nluc (Sec/SPI SP; predicted localization to OMV lumen) and BT_3630-Nluc (lipoprotein SP; predicted localization to OMV surface). PK, proteinase K (D) Schematic representation of post-secretion extracellular fate of Nluc mediated by BT_0169, BT_0569, and BT_3630 SP. BT_0922-Nluc is predicted to have a similar pattern as BT_3630 SP-Nluc as both have a lipoprotein SP and demonstrated similar abundance and localization in 7A and 7B. (E) Set of seven expression constructs generated to test the ability of BT_3630 SP to mediate secretion of different sized protein cargoes to the outer surface of OMVs in *B. theta*. The molecular weight of each protein is shown on the right. (F) Western blot analysis of liquid culture supernatants and cell pellets from *B. theta* expressing seven proteins of varying size fused to BT_3630 SP.

Next, we investigated the OMV-specific localization of two secretion carrier-Nluc fusions to further refine our ability to precisely implement therapeutic protein delivery with our engineered platform. We selected BT_0569 and BT_3630 SP as the secretion carriers for this study. Because BT_0569 has a Sec SP, it should be predominantly secreted into the periplasm through the Sec pathway. Conversely, BT_3630 SP is a lipoprotein SP which, in *E. coli,* can deliver protein cargoes to the inner leaflet of the OM through the localization of lipoprotein (Lol) pathway or the outer leaflet of the OM with additional secretion machinery^84^. Therefore, we predicted that BT_0569-Nluc should be packed into the OMV lumen and BT_3630 SP-Nluc should be anchored on the OMV membrane during the vesiculation process. We performed a proteinase K accessibility assay on OMV fractions isolated from liquid cultures of *B. theta* expressing BT_0569-Nluc and BT_3630 SP-Nluc **(Fig. 6C).** BT_0569-Nluc was highly resistant to degradation at both early (5 min) and late (30 min) timepoints across nearly all proteinase K concentrations tested, whereas BT_3630 SP-Nluc was much more sensitive to degradation at the higher proteinase K concentrations and over time. These results are consistent with our prediction and further imply that BT_3630 SP would anchor cargoes to the ‘outer leaflet’ of OMV membrane. Together these results suggest that the post-secretion fate (soluble, OMV surface, or OMV lumen) of protein cargoes can be controlled by careful selection of secretion carriers **(Fig. 6D),** which will allow more refined customization of *Bacteroides*-based *in situ* delivery systems for specific applications.

### Probing the size limit of lipoprotein SP-mediated protein secretion

To fully explore the capacity of *B. theta* for *in situ* delivery of protein-based therapeutics, we next wanted to determine if there is a limit on the size of the protein cargo that can be secreted by lipoprotein SPs with high functional secretion scores **(Fig. 3B).** We therefore selected BT_3630 SP as our representative SP and fused it to seven different cargo proteins – three endogenous *B. theta* proteins and four heterologous proteins – selected to cover a broad range of molecular weight: Nluc (25 kDa), cellulase (Cel; 41 kDa), BT_3686 (53 kDa), chitinase (ChiA; 68 kDa), BT_3703 (SusB; 88 kDa), β-galactosidase (LacZ; 122 kDa), and BT_3169 (148 kDa) **(Fig. 6E).** All constructs were fused with a C-terminal 3xFLAG tag to enable immunodetection. To minimize the metabolic burden and toxicity of constitutive high-level protein expression during the post-conjugation recovery and selection phase of growth, we used the aTc-inducible P1T_DP_-A21 promoter instead of the constitutive P_BfP1E6_-RBS8. Following growth in liquid culture to late log phase, we analyzed the culture supernatants for the presence of each secreted protein and only observed extracellular accumulation of the four smallest cargoes. The next largest protein cargo, SusB, was clearly observed in the pellet but not in the supernatant, suggesting an apparent molecular weight cutoff between 68 and 88 kDa **(Fig. 6F).** While it is possible that the observed low secretion levels of the two largest cargoes (LacZ and BT_3169) are a result of low expression levels, the results observed for SusB suggest the existence of a potential protein size limit for lipoprotein SP-mediated secretion.

### Secretion carriers mediate *in situ* delivery of heterologous proteins from *B. theta* in the mouse gut

Finally, to validate the *in vivo* functionality of our *in vitro*-characterized *B. theta* secretion carriers, we next investigated their performance in the gastrointestinal tract of mice. Following pre-treatment with an antibiotic cocktail, we orally gavaged C57BL/6J mice with: *B. theta* constitutively expressing Nluc with no secretion carrier (intracellular), *B. theta* constitutively expressing Nluc fused with BT_0294 SP (secreted; highest efficiency in secreting Nluc **(Fig. S1, 4C)**), wild-type (WT) *B. theta* (no heterologous protein secretion control), or PBS (no treatment control) **(Fig. 7A).** We monitored general health (mouse weight), *B. theta* colonization (colony forming units [CFU] in feces), and Nluc activity (luminescence) for two months **(Fig. 7A).** We observed no difference in weight between any group, suggesting that our engineered strains had no obvious adverse effects on mouse health **(Fig. 7B).** Both the intracellular and secreted Nluc strains engrafted and persisted at identical levels **(Fig. 7C),** demonstrating robust, long-term colonization despite competition with the native (antibiotic treated) mouse microbiota.

**Figure 7.**
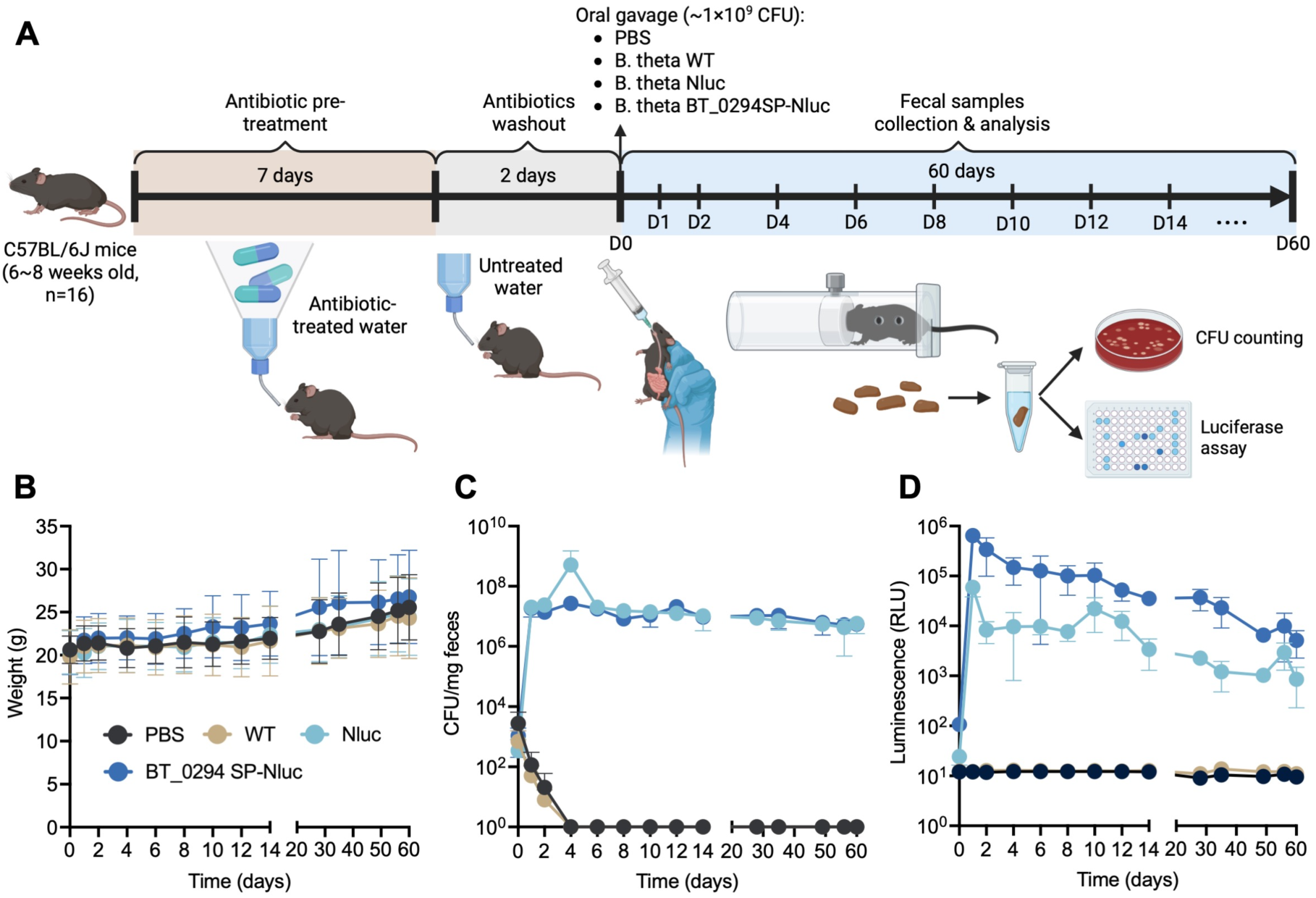
Direct intestinal delivery of heterologous protein cargo by *B. theta* in mice. (A) Design of *in vivo* experiments. Mice were monitored, and fecal samples were collected and analyzed for two months following inoculation. (B) The weight of mice in all groups increased similarly over time, indicating no adverse health effects. (C) Engineered *B. theta* strains persisted at high levels in the mouse intestine, as determined by fecal CFU counts. (D) The functionality of intestinally delivered protein cargo (Nluc) persisted over time, as determined by luminescence measurements of fecal homogenates. Error bars represent one standard deviation of the results of four mice for each group.

Because we performed these studies in antibiotic-treated conventional mice rather than germ-free, we used selective plating to isolate and quantify fecal CFU of our strains, which harbor the erythromycin-resistance gene. Thus, the CFU of the WT strain, which has no erythromycin-resistance marker, could not be quantified using this approach **(Fig. 7C).** Luminescence activity for both intracellular and secreted Nluc was readily detectable in the feces over the entire experimental timecourse, indicating that the secreted cargo was not only continuously present but also functional **(Fig. 7D).** The presence of Nluc in the fecal pellets of the mice colonized with the non-secreting (intracellularly expressing) *B. theta* strain suggests that some amount of cell lysis occurred *in vivo* in the mouse intestine or *ex vivo* during sample processing; however, the luminescence values measurements for fecal samples from mice colonized with *B. theta* expressing BT_0294 SP-Nluc (secreted) were around ten-fold higher than the intracellular Nluc values throughout the experimental timecourse. Despite stable *B. theta* colonization **(Fig. 7C),** we observed a slow decrease in luminescence activity over time for both the intracellular and secreted Nluc variants **(Fig. 7D).** While this is not entirely unexpected, as the expression constructs were extra-chromosomal and no antibiotic selection pressure was applied to maintain the plasmids in the *B. theta* cells, genome integration of expression/secretion constructs can be readily applied in future development and implementation of this approach.

## Discussion

*Bacteroides* species are a promising chassis for developing long-term interventions for diseases of the GI tract. However, the lack of efficient heterologous protein secretion tools for this genus limits their ability to serve as *in situ* production and delivery vehicles for therapeutic payloads or diagnostic reporters. In this work, we describe the development, characterization, and implementation of a molecular toolkit to enable efficient protein secretion in *Bacteroides* species. Previous studies attempting to address this need were able to achieve low levels of secretion of a number of different heterologous proteins, including human keratinocyte growth factor-2 (hKGF-2)^16,41^, murine interlukein-2 (mIL2)^39^, and human transforming growth factor β1 (hTGF-β1)^40^; however, secretion in these systems was mediated by Sec/SPI SP sequences from either *B. fragilis* enterotoxin^16,39,40^ or *B. theta* OmpA (BT_3852)^41^. Thus, similar to the results we observed with Sec/SPI SP secretion carriers, the secretion levels achieved in these studies were very low (<1 ng/mL). A key finding of our current work is the identification and characterization of a collection of lipoprotein SP secretion carriers and full-length protein secretion carriers derived from endogenous *B. theta* secretory proteins that enable high-titer secretion (up to 150 µg/mL) of functional heterologous proteins in multiple *Bacteroides* species. Interestingly, the two *E. coli* secretion systems we tested did not function at all in *B. theta*. This is likely due to the phylogenetic distance between these two species, which is much larger than the distance between *Salmonella enterica* serovar Typhimurium and *E. coli*^85^, for which successful secretion machinery swapping has been described^53^. Notably, several *Bacteroides* species are reported to have native T6SS and T9SS^37,38,86^, which are potential targets for future engineering and development of novel secretion tools.

Beyond our primary goals of achieving highly efficient protein secretion and identifying the sequence determinants that promote this optimized performance, we also undertook a detailed exploration of our engineered platform by examining specific behaviors of the system and its components. Toward the goal of developing a fully characterized and flexible platform for diverse biomedical applications, we identified a potential size limit of > 68 kDa and < 88 kDa as a cutoff for efficient secretion of heterologous protein from *Bacteroides* species. We also investigated the post-secretion extracellular fate of the heterologous protein cargo and identified differential secretion patterns depending on the secretion carrier. From these studies, we identified *B. theta* secretion carriers that appear to preferentially export protein cargo freely into the extracellular space (BT_0169), packaged within OMV lumen (BT_0569), or bound to the OMV surface (BT_3630 SP). Interestingly, BT_0922 seems to be cleaved near its C-terminus, leaving only a small portion still fused to the secreted heterologous cargo, making this a promising candidate for further development as a full-length secretion carrier. Notably, BT_0922 is conserved in many *Bacteroides* species, which could account for its high level of functionality across the *Bacteroides* species we tested, and further positions this secretion carrier as an attractive candidate for additional development.

Similar post-secretion protein localization results were recently reported for *E. coli* Nissle 1917 in which the SP from the *E. coli* lipoprotein Lpp, truncated OmpA, or hemolysin ClyA were fused to cargo proteins to direct them toward the lumen or external surface of OMVs^87^, suggesting that secretion machinery plays a key role in determining the localization of exported proteins on OMV. Notably, the Lpp SP promoted anchoring of protein cargo in the OMV membrane within the lumen, whereas the SP of the *B. theta* lipoprotein BT_3630 exported membrane-anchored proteins to the external OMV surface, while still other studies reported proteins targeted to both locations simultaneously when secreted by the lipoproteins P6 in *Haemophilus influenzae* or Pal in *E. coli*^88^. In another recent study also using *E. coli* as a model, cargo proteins fused with membrane proteins (OmpA, SLP, or SlyB), periplasmic proteins (BtuF or MBP), or a series of truncated Lpp SPs with different linkers demonstrated precisely controlled distribution and orientation in relation to OMVs^89^. Interestingly, the authors found that the size and number of OMVs produced by *E. coli* varied based on the fusion partner and fusion linker length, indicating a possible mechanism behind the cargo-dependent variability we observed with different secretion carriers in *Bacteroides* species. These features, together with the predicted size limit and post-secretion distribution that we observed, suggest that unknown lipoprotein transportation, sorting, and secretion systems exist in *Bacteroides*. Indeed, recent transposon mutagenesis studies in *B. theta* have revealed novel systems involved OMV production^90,91^, however, whether these systems are involved in lipoprotein sorting and protein packing into OMVs remains unclear and further investigation is required before these mechanisms can be exploited to improve secretion output.

The secretion carriers developed in this study enable enhanced heterologous protein secretion across multiple *Bacteroides* species. This platform expands the applicability of living therapeutics, which have previously focused largely on metabolic disorders and cancer in clinical development^4^ and have relied heavily on transiently colonizing probiotic strains, such as *E. coli* Nissle and *Lactococcus lactis*. By establishing a toolbox enabling the secretion of biotherapeutic proteins from permanently colonizing *Bacteroides* strains, we provide a means to utilize the living therapeutics platform for a broader range of diseases, including chronic conditions that require continuous treatment. Our additional characterization of the secretion carriers that we identified also provides a means for downstream users to select or engineer secretion carriers that are best suited for their particular goals and applications. Building on the success of our enhanced tunable promoter, future refinement of this platform will include incorporation of more sophisticated gene circuits for more precise control of expression/secretion outputs^23^, sense-and-respond circuits for disease-specific activation of therapeutic response^92–94^, and stable engraftment^26^ and biocontainment^25^ of engineered strains within the GI niche. Beyond therapeutic applications, *Bacteroides* species are prominent and abundant representative members of the gut microbiota^14^; the secretion tools described here could also be implemented for studying interspecies interactions and microbiota-host crosstalk in the gut.

## Acknowledgments

We thank S. Lyu for computational identification of lead peptide sequences. This work was funded by NIH NIBIB 1R21EB032548.

## Author contributions

Conceptualization: Y.H.Y. and S.J.S.; Methodology: Y.H.Y.; Formal Analysis: Y.H.Y.; Investigation: Y.H.Y. and V.W.K.; Resources: Y.H.Y., V.W.K. and R.R.P.; Writing – Original Draft: Y.H.Y. and S.J.S.; Writing – Review & Editing: Y.H.Y. and S.J.S.; Validation: Y.H.Y.; Visualization: Y.H.Y.; Supervision: Y.H.Y.; Project Administration: S.J.S.; Funding Acquisition: S.J.S.

## Methods

### Bacterial strains and culture

*Bacteroides thetaiotaomicron* VPI-5482, *Bacteroides fragilis* NCTC 9343, *Bacteroides ovatus* ATCC 8483, and *Bacteroides vulgatus* ATCC 8482 were acquired from ATCC. *Bacteroides* species were anaerobically cultured at 37 °C in TYG medium^23^, BHIS medium (Brain Heart Infusion Supplemented with 1 µg/ml menadione, 0.5 mg/ml cysteine, 0.2 mM histidine, 1.9 mM hematin) or on BHI agar with 10% horse blood (BHIB). *E. coli* strains were aerobically cultured in LB medium at 37 °C. *E. coli* DH5α was used for plasmid maintenance and *E. coli* RK231^95^ was used to achieve plasmid transfer in *Bacteroides* strains via tri-parental mating. Antibiotics were used when required at the following concentrations: ampicillin 100 µg/mL, kanamycin 50 µg/mL, gentamicin 25 µg/mL for liquid cultures and 200 µg/mL for agar plates, and erythromycin 12.5 µg/mL.

### Identification and selection of secretion carriers for fusion partner and OMV strategies

Based on the reported label-free quantification values (LFQ) of each *B. theta* protein in IM, OM, OMVp, or SUP fractions^43^, we calculated and ranked three different log_2_(ratio) values (log_2_[SUP/IM+OM+OMVp], log_2_[OMVp/IM+OM+SUP], and log_2_[SUP+OMVp/IM+OM]) for every detected *B. theta* protein to determine which proteins are highly secreted in the soluble fraction, insoluble OMV fraction, or both fractions (**Table S1**). From these three categories, we selected 57 secretory proteins (**Table S2**) to serve as secretion carriers for the fusion partner and OMV strategies. The 29 proteins in all three categories with a Sec/SPI SP and the single protein with a Tat/SPI SP were used as “full-length” secretion carriers in the fusion partner strategy based on the assumption that their extracellular export relies on protein domains other than just their SPs. In addition, 4 lipoproteins with high log_2_(SUP/IM+OM+OMVp) values were also used as “full-length” secretion carriers in the fusion partner strategy based on previous reports in *E. coli*^84^ demonstrating that lipoprotein SPs are only responsible for anchoring proteins to the IM or OM, and other components of the secreted protein may help mediate extracellular export. For the remaining 23 lipoproteins with high log_2_[OMVp/IM+OM+SUP] or log_2_[SUP+OMVp/IM+OM], we identified the N-terminal charged and central hydrophobic regions preceding the cysteine cleavage site and used this along with ∼20 additional amino acids following the cleavage site as the secretion carriers for the OMV strategy. BT_p548220 is a lipoprotein expressed from an endogenous *B. theta* plasmid and has a high log_2_(SUP+OMVp/IM+OM); it was selected as part of the OMV strategy due to its homology with a highly secreted *B. fragilis* plasmid-expressed protein, BF9343.20c^96^. BT_2472, though it has the highest log_2_(SUP/IM+OM+OMVp), was not selected because we could not detect its secretion after C-terminal fusion with 3xFLAG (Fig. 1).

### Molecular cloning

Q5 high-fidelity DNA polymerase (New England Biolabs) was used for PCR amplification of DNA fragments for cloning. All primers were synthesized by Integrated DNA Technologies (IDT). All plasmid construction was done by Gibson Assembly (HiFi DNA Assembly Master Mix, New England Biolabs) and validated by colony PCR and sequencing. Plasmids were stored in *E. coli* DH5α for maintenance and conjugation. To express proteins in *Bacteroides* species, we constructed a *E. coli*-*Bacteroides* shuttle vector, pBT (**Fig. S2**), by assembling the *repA* (amplified from pAFD1^97^; responsible for plasmid self-replication in *Bacteroides* species), *traJ*/RK2 *oriT* (amplified from pAFD1; responsible for the initiation of plasmid conjugal transfer), *mobA* (amplified from pFD340; assist plasmid transfer during conjugation), *ermG* (erythromycin resistance; amplified from pNBU2), *ampR*, and pBR322 *ori* via Gibson assembly. All endogenous secretion carriers were cloned from the genome of *B. theta*. The HlyA, HlyB, HlyD of UPEC T1SS were cloned from pVDL9.3 (Addgene #168299) and pEHlyA5 (Addgene #168298) plasmids. The CsgG of *E. coli* K-12 T8SS was cloned from the genome of *E. coli* DH5α. For constructs with N-terminal fusions (CsgA [N-terminal 22 residues], SusB signal peptide, and BT_3769 signal peptide), the sequences were introduced at the N-terminus of sdAb- TcdA directly through primers. The sequences of P_BfP1E6_, sdAb-TcdA, VHH3, and EGFP were synthesized by IDT. The sequences of Nluc and 7D12 were cloned from plasmids pNBU2_erm-TetR-P1T_DP-GH023-NanoLuc (Addgene #117728) and pTrcHIS-wt7D12 (Addgene #125268). The scFv-HER2 was constructed from trastuzumab as previously described^68^. All the promoters, RBS, secretion carriers, and protein cargoes were cloned into pBT vector at the sequence between *ampR* and *ermG*, and 3xFLAG as well as a rrnb T1 terminator of *E. coli* was further introduced at the end of the protein cargo coding region by overhang primer PCR. A GSSG or GSSGSSG linker was introduced at the C-terminus of each protein, in frame with the 3xFLAG tag and a GSGGSGSSGS linker was introduced at C-terminus of each full-length secretion carrier, in frame with the protein cargo, by overhang primer PCR. For the plasmids used for recombinant protein expression and purification, the toxin A fragment (TcdAf; amino acid residues 2460-2710) was amplified from the *C. difficle* genome by PCR and cloned into the 2Bc-T plasmid (Addgene #37236); Nluc, sdAb-TcdA, sdAb-TNFα, and sdAb-EGFR were cloned into the pET24b(+) plasmid with a 3xFLAG tag at the C-terminus and an N-terminal fusion with the *E. coli* outer membrane protein A (OmpA) signal peptide to for secretion of proteins into the periplasm^98^.

### Conjugation and selection

Plasmids for *Bacteroides* conjugations were first used to transform *E. coli* DH5α to generate plasmid donor *E. coli* strains. Overnight cultures of plasmid donors, *E. coli* RK231 (helper strain), and *Bacteroides* (plasmid recipient) were combined in a 1:1:1 ratio of volume. The mixed liquid cultures were pelleted by centrifugation at 10,000 xg for 1 min, resuspended in 30 µL LB medium, spotted onto on BHIB plates, and incubated aerobically for 24 hr at 37 °C. Mating spots were scraped off plates, streaked onto BHIB plates supplemented with selective antibiotics (200 µg/mL gentamicin and 25 µg/mL erythromycin), and incubated anaerobically for 2-3 days at 37 °C to allow selective growth of transconjugant *Bacteroides* clones.

### Recombinant protein expression and purification

The HER2 extracellular domain (ECD) was purified as previously described^68^. For sdAb-TcdA and toxin A fragment (TcdAf) purification, an overnight culture of *E. coli* BL21(DE3) harboring the 2Bc-T-TcdAf plasmid was grown overnight at 37 °C with shaking, then diluted 50-fold in 50 mL terrific broth (yeast extract 24g, tryptone 20g, glycerol 4mL, 100 mL KPO_4_ buffer [0.17 M KH_2_PO_4_, 0.72 M K_2_HPO_4_] per liter of medium) with 50 µg/ml kanamycin. When culture OD_600_ reached 0.6, isopropyl β-D-thiogalactoside (IPTG) was added to a final concentration of 0.1 mM to induce protein expression. After overnight induction of cultures at 25 °C with shaking, the cells were harvested and sonicated in lysis buffer (20 mM sodium phosphate, 0.5 M NaCl, 40 mM imidazole, 1% Triton X100, 0.1 mM PMSF pH 7.4). The soluble fractions of cell lysates were passed through a Ni-NTA chromatography column, and the recombinant TcdAf proteins were eluted with elution buffer (20 mM sodium phosphate, 0.5 M NaCl, and 500mM imidazole). The concentration of purified TcdAf was calculated from A280, molecular weight, and extinction coefficient based on Beer-Lambert law. For Nluc, sdAb-TcdA, sdAb-TNFα, and sdAb-EGF, the expression plasmids were introduced into *E. coli* BL21(DE3). Overnight cultures of *E. coli* BL21(DE3) harboring these plasmids were diluted 50-fold in 100 mL BHI with 50 µg/ml kanamycin. After the OD_600_ reached 0.3, protein expression was induced with 0.1mM IPTG and cultures were grown at 27 °C for sdAb-TcdA, sdAb-TNFα, and sdAb-EGFR and at 22 °C for Nluc, with shaking, overnight. The cells were pelleted by centrifugation at 5000 xg for 10 min and sonicated in lysis buffer (PBS with 1% Triton X-100 and 0.1 mM PMSF). The soluble fraction of cell lysates was loaded to a column with anti-FLAG® M2 affinity gel (Sigma-Aldrich). Because 3xFLAG tagged scFv-HER2 did not express well in *E. coli* BL21(DE3), we purified it from the supernatant of a culture of *B. theta* expressing BT_3630SP-scFv-HER2-3xFLAG, loaded directly onto the anti-FLAG column. The column was washed once with PBS and the protein was eluted with 0.1 M glycine (pH 3.5). The protein concentration was determined by DC protein assay (Bio-Rad).

### Measurement of protein secretion by dot blot or western blot analysis

*Bacteroides* strains were first streaked on BHIB plates with antibiotics (200 µg/mL gentamicin and 25 µg/mL erythromycin). Colonies were inoculated into TYG or BHIS media with 12.5 µg/mL erythromycin (with 100 ng/mL aTc when using aTc-inducible promoters). *Bacteroides* strains harboring plasmids with constitutive promoters were grown to stationary phase while those with aTc-inducible promoters were grown to early log phase. The culture supernatants were separated from bacterial cells by centrifugation at 10,000 xg for one minute and filtered through 0.22 µm PVDF syringe filters. For dot blot analysis, 10-30 µL of filtered supernatant was directly spotted onto a wet PVDF membrane placed on a stack of paper towels to aid rapid and consistent wicking of the samples through the membrane. For western blot analysis, 10.5 µL of filtered supernatant was mixed with 1.5 µL β-mercaptoethanol and 3 µL 5x sample loading buffer (300 mM Tris-HCl at pH 6.8, 10% SDS, 50% glycerol, and 0.5% bromophenol blue dye) and boiled at 98°C for 10 min. 10µL of boiled sample was subjected to Tris-glycine SDS-PAGE (12% gel or 4-20% gradient gel) in electrophoresis buffer (25 mM Tris-base, 250 mM glycine, and 0.1% SDS) at 100V for 1.5 hr for protein separation. Proteins were transferred to a PVDF membrane at 100V for 1 hr in transfer buffer (25 mM Tris-base, 192 mM glycine, and 30% methanol). For both dot blots and westerns, the membranes were blocked with 5% milk in 0.1% PBS-T (phosphate-buffered saline with 0.1% Tween 20) at room temperature for 1 hr, then incubated with anti-FLAG M2 monoclonal antibody (Sigma-Aldrich, 1:2000 dilution in 5% milk) at 4 °C with rocking overnight. After washing three times with PBS-T, the membrane was incubated with goat anti-mouse IgG secondary antibody conjugated with horse radish peroxidase (HRP) (Jackson Immuno Research, 1:5000 dilution in 5% milk/0.1% PBS-T) at room temperature for 1 hr. Signal was detected using SuperSignal™ West Dura Extended Duration Substrate (Thermo Scientific) on a Bio-Rad GelDoc imaging system. For dot blot, the signal intensity was quantified by ImageJ. Briefly, the color of dot blot image was first inverted to obtain a black background, and the signal intensity of each dot was determined by calculating the mean gray value in the circular selection area via ROI Manager.

### Measurement of activities of secreted antibody fragments and reporters in culture supernatants

“Activity” is defined as antigen binding for antibody fragments and enzymatic or fluorescent activity for reporter enzymes. The activities of all antibody fragments were measured by ELISA as follows: 96-well Immulon 2HB ELISA plates were coated with 2 µg/ml purified antigens (TcdAf/HER2 ECD, purified as described above; TNFα soluble form (#10602-HNAE-100) and EGFR ECD (#10001-H08H) were purchased from Sino Biological) at 4 °C overnight. After washing 3x with 0.1% PBS-T, plates were blocked with 5% milk/0.1% PBS-T for 1 hr at room temperature and washed again. Filtered culture supernatants (100 µL) were added to wells and incubated for 1hr at RT. Plates were washed again and anti-FLAG M2 antibody (Sigma-Aldrich, 1:2000 dilution in 5% milk/0.1% PBS-T) was added and incubated for 1hr at room temperature. Following PBS-T washing, goat anti-mouse IgG secondary antibody conjugated with HRP (Jackson Immuno Research, 1:5000 dilution in 5% milk/0.1% PBS-T) was added and incubated for 1hr at RT. After washing with PBS-T, o-phenylenediamine (OPD) substrate solution was added and allowed to react for 30 min at room temperature. The absorbance at 450 nm (A450) was measured using a BioTek Synergy HT multimode microplate reader. To quantify secreted sdAb-TcdA, sdAb-TNFα, sdAb-EGFR, and scFv-HER2, 10-fold serially diluted purified 3xFLAG tagged proteins were run in parallel as standards, and the four-parameter logistic (4PL) regression model was applied to build the sigmoidal standard curves to estimate the secretion titer. For NanoLuc, the Nano-Glo luciferase assay (Promega) was performed as follows: 10 µL filtered culture supernatant was mixed with 15 µL PBS, followed by mixing with 25 µL Nano-Glo® luciferase assay buffer supplemented with furimazine substrate at a ratio of 1:50. Luminescence was measured on the microplate reader using an integration time of 1 s and gain of 100. Quantification of secreted Nluc was achieved with a 10-fold serial dilution of purified 3xFLAG tagged Nluc and a standard linear regression model was applied to build the standard curve for quantifying secreted Nluc. For β-lactamase, the β-lactamase activity assay kit (Sigma-Aldrich) was used following the manufacturer’s protocol. Briefly, 10 µL filtered culture supernatant was mixed with 40 µL PBS, followed by mixing with 50 µL β-lactamase assay buffer supplemented with nitrocefin substrate at a ratio of 1:25. Plates were incubated for 5 min at room temperature, then the absorbance at 490 nm (A490) was measured using the microplate reader.

### Analysis of the post-secretion extracellular fate of secreted proteins

*B. theta* colonies were inoculated into 30 mL TYG with 12.5 µg/mL erythromycin and anaerobically grown to late log phase or stationary phase. 1 mL of liquid culture was first centrifuged at 10000 xg for 1 min to separate cell pellets and culture supernatant. Cell pellets were then washed once with PBS to obtain the “cell pellet” fraction, which was then resuspended in 200 µL 2x sample loading buffer with 10% β-mercaptoethanol, boiled at 98 °C for 10 min, and diluted with 200 µL PBS to bring the sample buffer to 1x. The remaining 29 mL of culture was centrifuged at 5000 xg for 10 min to separate the cells and supernatant, and the supernatant was filtered through a 0.22 µm syringe filter to remove remaining cells and debris to obtain the cell-free “total supernatant” fraction. OMVs in the total supernatant fraction were extracted using the ExoBacteria™ OMV Isolation Kit (System Biosciences) according to the manufacturer’s protocol: 25 mL of the total supernatant fraction was transferred to an OMV-binding column and rotated at 4 °C for 30 min. The column was uncapped, and the flow through was collected as the OMV-free “soluble” fraction; finally, the column was washed twice with 15 mL binding buffer, and the OMVs were eluted with 1.25 mL elution buffer to obtain the 20x-concentrated “OMV” fraction. For western blot analysis, 10 µL from the total supernatant, soluble, and OMV fractions were mixed with 1.5 µL β-mercaptoethanol and 3 µL 5x sample loading buffer and boiled at 98 °C for 10 min. SDS-PAGE and immunoblotting were performed as described above with 10 µL of each processed fraction. For Nluc quantification, 10 µL of total supernatant, soluble, and OMV fractions containing secreted Nluc were analyzed using the Nano-Glo assay as described above.

### Animal experiments

All animal experiments were performed using protocols approved by the University of Illinois Institutional Animal Care and Use Committee. C57BL/6J specific-pathogen-free (SPF) mice (6-8 weeks old; sex-balanced) were pre-treated with an antibiotic cocktail in their drinking water (1 g/L metronidazole, 1 g/L neomycin, 0.5 g/L vancomycin, and 1 g/L ampicillin, and 20 g/L Kool-Aid Drink Mix^99^) for 7 days followed by a 2-day washout period with plain tap water. Mice were divided into four groups (2 males and 2 females per group) and administered 200 µL of PBS or bacterial strains (1×10^9^ CFU). Mice were weighed and fecal samples were collected every two days for two weeks, then once per week for another six weeks, with a final sample collected on day 60. To quantify colonization of the engineered *B. theta* strains, fecal samples were homogenized in PBS (10 µL per mg of feces) and serially diluted in 96-well plates. Dilutions were plated on selective BHIB plates using a multi-channel pipettor and the highest dilution with single colonies was used to calculate CFU per mg of feces. To measure the level of secreted Nluc in fecal pellets, the homogenized fecal pellets were centrifuged at 12000 xg for 2 min to pellet bacterial cells, then 25 µL of supernatant was used in the Nano-Glo luciferase assay described above.

## Supplmental Figures and Tables

**Figure S1.**
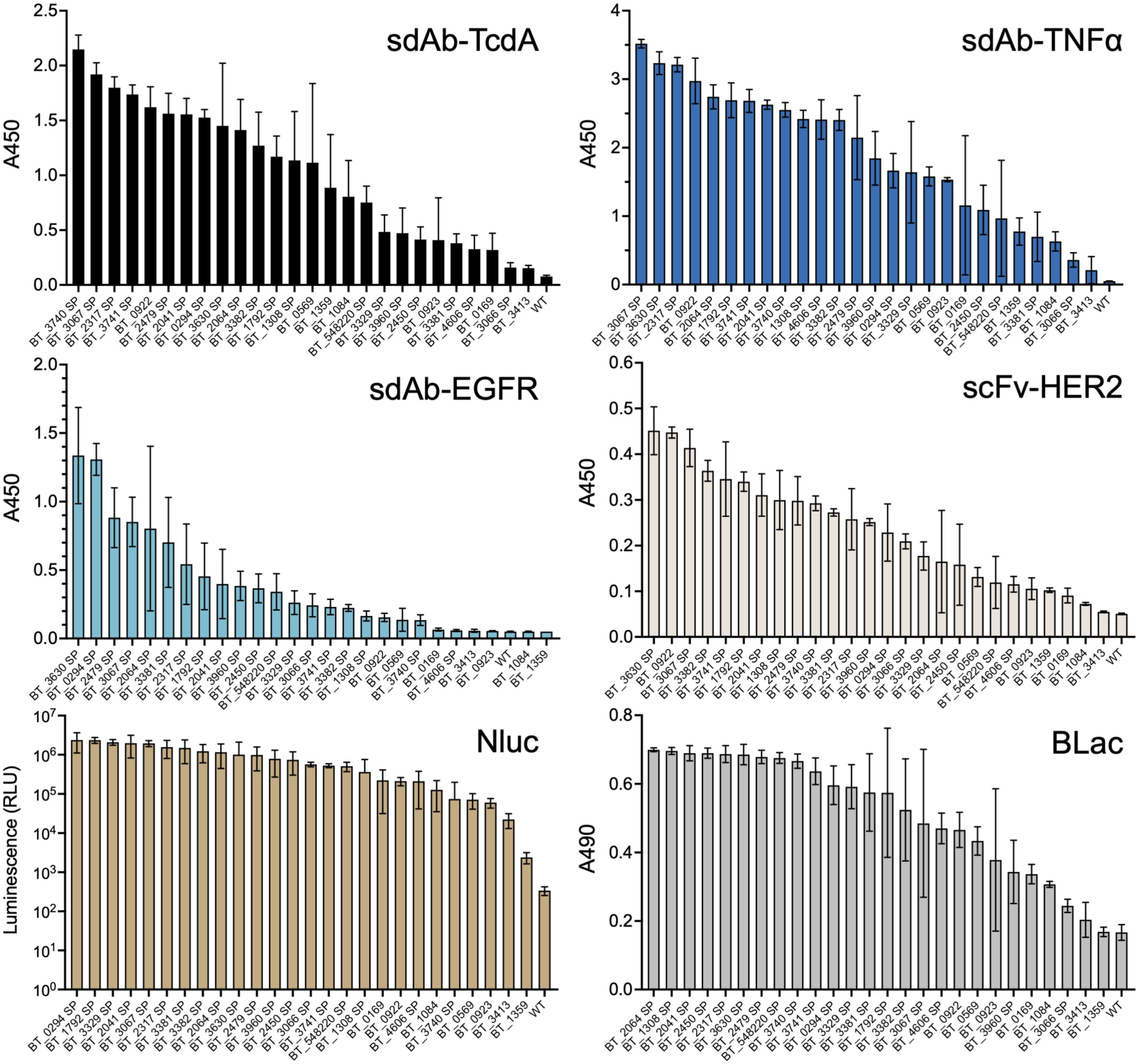
Activity assays of antibody fragments and reporter proteins secreted from B. theta. The activities of antibody fragments (sdAb-TcdA, sdAb-TNFα, sdAb-EGFR, and scFv-HER2) and reporter proteins (Nluc and BLac) secreted by 26 secretion carriers in B. theta culture supernatants were measured by functional assays (ELISA for antibody fragments; luciferase assay for Nluc; colorimetric enzymatic assay for BLac). The secretion carriers were ranked based on their average readouts in triplicate experiments. Error bars represent one standard deviation of triplicate biological samples.

**Figure S2.**
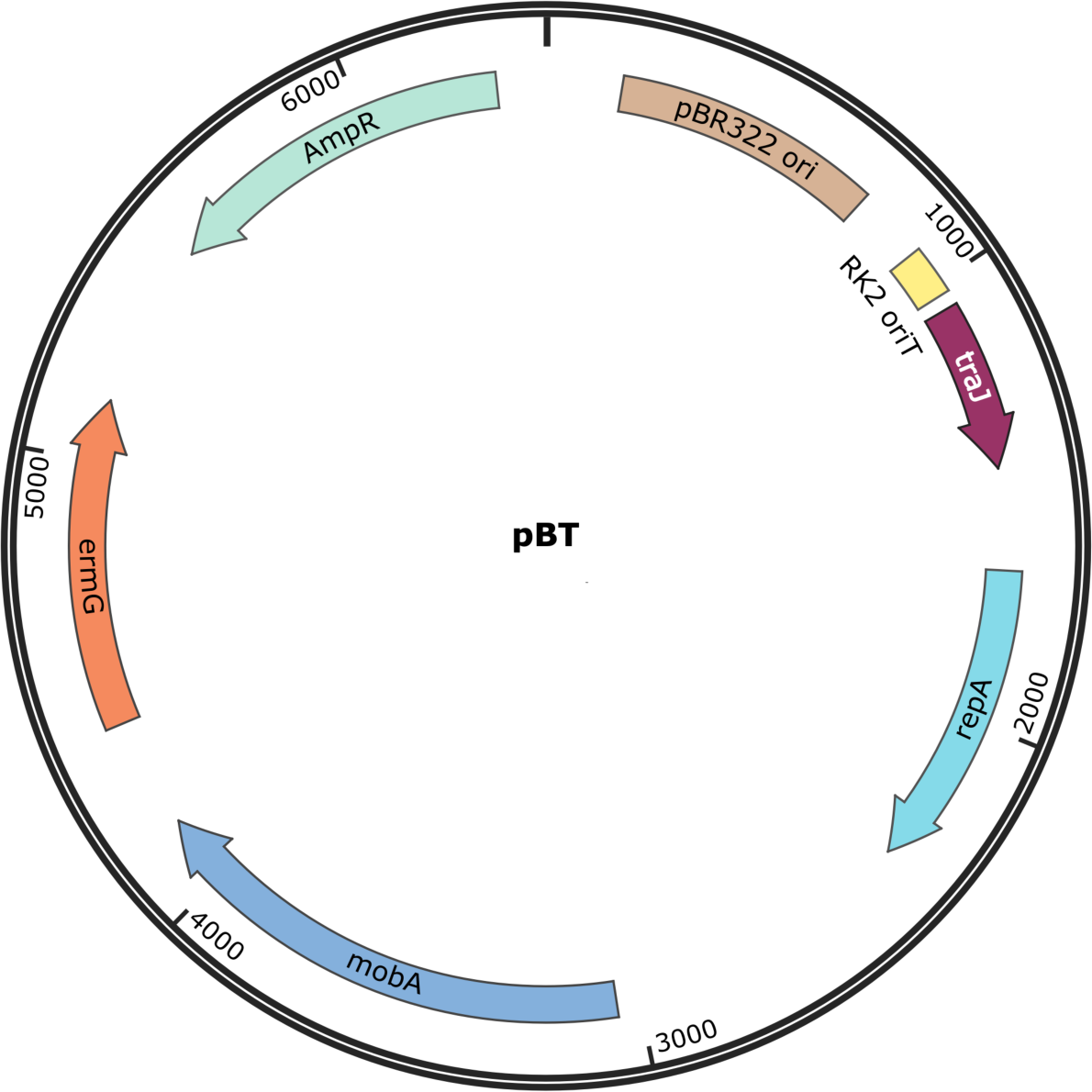
Vector map of pBT. The plasmid map was created by SnapGene Viewer.

**Table S3.** Solubility and pI of seven protein cargoes.

|  | Predicted scaled solubility<sup>a</sup> | pI<sup>a</sup> |
| --- | --- | --- |
| sdAb-TcdA | 0.503 | 9.06 |
| sdAb-TNFα | 0.429 | 5.9 |
| sdAb-EGFR | 0.516 | 9.08 |
| scFv-HER2 | 0.457 | 9.3 |
| EGFP | 0.607 | 5.76 |
| Nluc | 0.581 | 5.12 |
| BLac | 0.454 | 5.53 |

<sup>a</sup>The solubility and pI were predicted and calculated by Protein-Sol (https://protein-sol.manchester.ac.uk)

